# Sex and morph variation in activity from early ontogeny to maturity in ruffs (*Calidris pugnax*)

**DOI:** 10.1101/2024.08.30.610446

**Authors:** Veronika A. Rohr-Bender, Krisztina Kupán, Guadalupe Lopez-Nava, Wolfgang Forstmeier, Anne Hertel, Vitali Razumov, Katrin Martin, Bart Kempenaers, Clemens Küpper

## Abstract

Intraspecific variation provides the substrate for the evolution of organisms. Ruffs show exceptional phenotypic variation in physiology, appearance and behaviour linked to variation between sexes and male alternative reproductive tactics (ARTs). The male ARTs are associated with the evolution of separate morphs, which are encoded by an autosomal supergene. However, the effects of the supergene on females and chicks are much less well understood. In particular, it is still unknown, whether females also show morph-specific behavioural variation, when behavioural differences emerge during ontogeny, and whether behavioural differences can be detected outside of the breeding context. To address these knowledge gaps, we repeatedly measured the activity in an unfamiliar environment, also known as exploration behaviour, of 109 hand-raised young ruffs throughout their first two years of life. We used automated tracking in an open field arena, and quantified the distance moved within 10 minutes to examine behavioural differences between sexes, morphs and individuals. The activity of young ruffs rapidly increased during the first month after their crouching reflex, a response to potential threats, subsided. Repeatability of individual activity was initially low but increased throughout juvenile ontogeny and was high (R = 0.5) from day 21 onwards. Variation in activity was clearly sex-linked with females moving more than males, indicating potential energetic trade-offs accompanying the strong sexual size dimorphism. In contrast, morph differences in activity remained inconsistent and elusive, both in females and in males. Our results indicate that in species where much of the known behavioural variation is linked to mating tactics, a non-reproductive behaviour can show between-individual variation and clear sex differences, whereas morph differences appear less pronounced.

## Introduction

An individual’s activity in a novel environment is a well-studied personality trait and is commonly referred to as exploration behaviour (Dingemanse et al. 2002; Réale et al. 2007). This activity has been related to foraging behaviour (Herborn et al. 2010; Ersoy et al. 2022), home range size (Boyer et al. 2010; Minderman et al. 2010), large-scale patterns of space use (Bijleveld et al. 2014) and dispersal (Cote et al. 2010), as well as to reproduction (Schuett et al. 2012; Mutzel et al. 2013) and survival (Dingemanse et al. 2004; Smith and Blumstein 2008; Bergeron et al. 2013). Additionally, individual variation in activity has been linked to hormone levels, such as testosterone (van Oers et al. 2011; Raynaud and Schradin 2014). Activity is commonly assessed using open field tests, which involves placing the animal in a novel, open arena and measuring its movements and interactions with the environment (Hall and Ballachey 1932; Perals et al. 2017). Repeated measurements of individuals allow then the assessment of between-individual differences in behavioural means via variance standardization (i.e., repeatability, Bell et al. 2009).

Personality traits are typically measured in adult animals, while few studies have documented the development of between-individual behavioural variation during early juvenile ontogeny (Bell et al. 2009; Kok et al. 2019; Ersoy et al. 2024). The few existing studies conducted during early ontogeny revealed that individual behavioural repeatability typically increases during development, as between-individual variation increases and within-individual variation decreases with age (Fisher et al. 2015; Carlson and Tetzlaff 2020; Freund et al. 2013; Polverino et al. 2016; but see Bell et al. 2009; Kok et al. 2019). However, it remains unclear when exactly between-individual differences solidify.

The study of between-individual differences in behaviours involving movement during early ontogeny poses several challenges. First, locomotive abilities increase with age during development due to physical growth, resulting in a significant increase in speed, often by orders of magnitude (Hertel et al. 2023). These strong developmental changes might hinder the detection of smaller effects, such as between-individual differences. Second, the mode of locomotion may change during development. Shifts in locomotor type, such as transitioning from swimming to walking in amphibians or from walking to flying in birds and insects, can interfere with experimental setups that are designed to test the same behavioural trait. Third, the consideration of early ontogeny up to adulthood offers a comprehensive perspective, but this period can be quite long, especially in long-lived species. This may affect repeatability estimates, as repeatability likely decreases with increasing length of the test period similarly to the length of the interval between observations. This is because within-individual variance tends to be higher when measured over a longer time period and there is more opportunity for developmental change (Bell et al. 2009; Holtmann et al. 2017). Fourth, variance components are often not reported. Variance components underlying repeatability, i.e., within- and between-individual variation, can increase, decrease or remain constant independently of each other and frequently are not evaluated or reported (Bell et al. 2009; Holtmann et al. 2017; Dingemanse et al. 2022).

Investigating between-individual variation in behaviours in juveniles goes beyond a purely mechanistic understanding of the development of animal personality. The juvenile stage is a critical life period, where between-individual variation in activity-related behaviours may come with benefits or reflect important trade-offs that are relevant to fitness. First, activity influences predation risk. High levels of activity may reflect high vigour, as it has been associated with better maximal locomotor performance than low levels, increasing the likelihood of escaping predators (Careau and Garland 2012) and ultimately survival (Smith and Blumstein 2008). For example, yearling brown trout (*Salmo trutta*) exhibiting higher velocity in an open field test were more likely to survive during the initial months of their lives (Adriaenssens and Johnsson 2013). Conversely, activity may increase the risk of encountering or being detected by predators, resulting in a trade-off between activity and survival (Lima and Dill 1990). For example, juvenile European rabbits (*Oryctolagus cuniculus*) with higher exploration scores had lower survival probabilities than those with lower exploration scores (Rödel et al. 2015). Second, activity is linked to foraging and growth. Proactive individuals often engage in more active and risky foraging behaviours, which can facilitate faster growth (Réale et al. 2007; Hussey et al. 2017). However, due to a trade-off between activity and growth, allocating a significant amount of energy to activity-related behaviours may limit the resources available for physiological processes such as body maintenance and growth, which may be detrimental to survival when resources are limited (Careau et al. 2008).

In addition to age and between-individual differences, behavioural variation can be explained by categorical differences, such as sex or morph. Within many species, the two sexes are a major source of activity-related behavioural variation. Males and females often differ for example in territory size (Wikramanayake and Shrestha 2014), and in breeding and natal dispersal (Greenwood 1980; Pusey 1987). Reproductive morphs that pursue alternative reproductive tactics (ARTs) are another source of behavioural differences in the context of mating and reproduction (Oliveira et al. 2008). Some of the differences between ARTs go hand in hand with differences in activity. For example, in both sexes of prairie voles (*Microtus ochrogaster*), a species with two alternative morphs, socially monogamous residents establish pair bonds and territories, whereas polygamous wanderers remain non-pair bonded, non-territorial and generally move greater distances (McGuire and Getz 2010). Behavioural differences between sexes or morphs are usually most pronounced in adults, but may already emerge during early ontogeny (Lynn and Brown 2009; Krzyszczyk et al. 2017). However, exactly when during ontogeny these differences between sexes and morphs arise is often unclear.

We used an open field test paradigm to investigate the development of sex-dependent, morph-dependent and individual variation in activity in young ruffs (*Calidris pugnax*). The ruff sandpiper is exceptionally suitable for studying this for several reasons. Beside the differences between sexes there is substantial variation within the sexes (Hogan-Warburg 1966; van Rhijn 1991; Widemo 1998; Lank et al. 1995; Lank et al. 2013; Jukema and Piersma 2006). Ruffs can be assigned to one of three morphs. These morphs are encoded by an autosomal supergene, which means that the three genetic morphs are present in both males and females (Lamichhaney et al. 2016; Küpper et al. 2016). During breeding season in males the morphs manifest as different male ARTs in this lekking species: (a) The majority of males are “Independents”, who aggressively defend territories on leks; (b) Fewer males are “Satellites”, who take the role of a subdominant partner in a courtship alliance with an Independent; (c) The rarest morph are “Faeders”, who exhibit female-like plumage and behaviour to sneak copulations. However, to date, no behavioural differences between the female ruff morphs have been described. Additionally, very little is known about how the genetic differences may affect general fitness or behaviour outside of the reproductive context and during early life. Further, the three morphs differ in circulating hormone concentrations (*inter alia* testosterone) not only in adult males (Küpper et al. 2016; Loveland et al. 2021) but in both sexes already at the chick stage (Giraldo-Deck et al. 2022). Independents have higher plasma testosterone concentrations than the other two morphs and these endocrinal differences may influence activity, given that testosterone concentrations often correlate with activity (van Oers et al. 2011; Raynaud and Schradin 2014). Additionally, ruffs show striking intraspecific variation in body size. Adult males are approx. 70 % larger than females (Münster 1990; Meissner and Zięcik 2005). Within both sexes, the three morphs differ in size, with Independents being the largest, Satellites intermediate and Faeders being the smallest (Lank et al. 2013; Giraldo-Deck et al. 2020). These size differences already emerge during the first week of life (Giraldo-Deck et al. 2020) and could be accompanied by behavioural differences. Ruff chicks are precocial, become active and leave the nest within a few hours of hatching. During their first weeks of life, wader chicks face a high mortality risk due to predation and must soon make increasingly independent decisions from their parents (summarised by Colwell et al. 2007). Sex and morph variation in activity may alter predation risk and differently affect survival prospects of certain morphs or sexes.

To investigate the development of activity, we conducted a series of open field tests. We tested captively raised ruffs six times during juvenile ontogeny until physical growth was mostly complete (<33 days old), and again three times during their first and second winters (approx. ½ and 1 ½ years old, respectively). The winter tests allowed us to examine the behavioural stability of their mature phenotypes. Activity, as measured here, is traditionally referred to as exploration behaviour, which refers to an individual’s movements in a new environment (Réale et al. 2007; Dingemanse et al. 2022). However, recently, concerns have been raised about the decline of novelty due to repeated testing and the interpretability of this behaviour as being exploratory (e.g., Dingemanse et al. 2002; Minderman et al. 2010; Dingemanse et al. 2012; Carter et al. 2013; Bijleveld et al. 2014; Wuerz and Krüger 2015; Perals et al. 2017). To avoid a biological interpretation of this behaviour as being exploratory, we therefore refer to it as ‘activity in an unfamiliar environment’ (in short, ‘activity’). We predicted an increase in activity in young ruffs over time until a mature phenotype is reached in the birds’ first winter. Due to the prevalent differences in size, physiology and reproductive behaviour, we investigated whether activity was related to an individual’s sex and morph. Given the limited knowledge about sex- and morph-specific differences in the development of activity, our approach remains exploratory. Our main prediction is that sex will have a stronger effect on the young ruffs’ activity than morph, because size and behavioural differences in adults are larger between the sexes than between the morphs. We had no clear predictions if potential morph differences should vary by sex, therefore, we refrained from exploring a potential interacting effect between sex and morph. To investigate individual variation, we calculated the between-individual repeatability of activity and assessed how it changes across different life stages. We predicted an increase of repeatability with age until the mature behavioural phenotype is reached from the first winter onwards.

## Methods

### Study population

We conducted this study using a captive population of ruffs at the Max Planck Institute for Biological Intelligence in Seewiesen, Germany over three consecutive breeding seasons (2021-2023). The founder individuals of this population were provided by David Lank who raised ruffs from eggs collected near Oulu, Finland, in 1985, 1989, and 1990 (Lank et al. 2013) and maintained a captive population many years at Simon Fraser University in Burnaby, British Columbia, Canada, before the birds were translocated to Seewiesen in 2019. We supplemented the population with further adult ruffs obtained from ruff breeders and zoological gardens in the Netherlands, Belgium and Germany. We incubated eggs artificially, and hand-raised chicks on *ad libitum* food. After hatching, we collected a blood sample (5-10μl) to determine sex and morph using diagnostic single nucleotide polymorphism markers (see Giraldo-Deck et al. 2020). After hatching, we kept the chicks in small groups of up to 10 individuals in plastic boxes (60 × 40 × 32 cm) for the first 9-10 days of their lives. Subsequently, we transferred them to larger indoor aviaries (2 × 3 m) housing up to 35 juveniles. At 33 days of age, we introduced the juveniles to the main flock of adult birds in a large aviary (∼150 m^2^).

### Open field test

We designed an open field test to assess individual behavioural responses in a controlled, unfamiliar environment. The experimental arena (2 × 2 m) was a plain, smooth floor marked with a 4 × 4 field grid using 10 mm wide tape, bordered by the white walls of the test room. We tested each individual separately, directly transferring it from the housing environment into the experimental room. Before each trial, we placed the bird under an upside-down black flower pot. At the start of each trial, we raised the pot from outside the room via a wire rope hoist and recorded the individual’s movements by a top-view camera for a duration of 10 minutes. The experimental arena presented a novel environment during the initial exposure, and an unfamiliar environment in subsequent repeats.

To capture the development of activity, we tested the chicks every 6 days, from day 3 to day 33 post-hatching, resulting in up to six open field tests during their juvenile ontogeny. The first three trials were carried out during the early juvenile ontogeny, prior to fledging. The subsequent three trials took place during later juvenile ontogeny (**Fehler! Verweisquelle konnte nicht gefunden werden**.), extending to almost the end of the physical growth phase (Giraldo-Deck et al. 2020). The testing order was random within the age groups but opportunistic between age groups, i.e., depending on other behavioural experiments scheduled on a given day. For 3% of the scheduled trials, we shifted individual tests by one day. Chicks that were temporary unwell (e.g., had substantially lost weight compared to the last day, did not show a flight response when approached) on a test day were not tested meaning that 24% of the 271 test individuals who survived until day 33 did not undergo all six open field tests (see **Fehler! Verweisquelle konnte nicht gefunden werden**. for details). To assess the mature phenotype of activity, we tested the juvenile ruffs three times with a 6- and 7-day interval during the second half of November and early December in their first winter (approx. ½ year old; 24^th^ Nov - 9^th^ Dec in 2021, 15^th^ Nov - 29^th^ Nov in 2022, and 17^th^ Nov - 30^th^ Nov in 2023). During each trial, the random order in which we captured them from their home compartment determined the testing order. We tested all young ruffs that remained at our facility in their second winter (approx. 1½ years old) on the same designated test days. Due to a technical problem with video recordings, we repeated the third trial for 14 of 124 individuals tested (first winter test for 11 individuals, second winter test for 3 individuals) in November 2022 together with 10 additional individuals that served as a control group on the next day. The activity measured in the control group was highly correlated between the repeated and original third trial (Pearson’s correlation, r = 0.68, n = 10) and consequently we included the repeated tests of all 24 individuals for all further analyses.

### Automated Tracking & Track Cleaning

As a measure of activity, we quantified the distance travelled by birds in each open field test using automated tracking from 109 individuals (Table 1, Figure S1). We aimed for a balanced representation of all sex-morph classes, but we were limited by the number of individuals from the rare Faeder morph and Satellite males that survived juvenile ontogeny. The number of males and females in the analysed sample was overall similar (±10 %). Where possible, we selected individuals that had completed all trials during the juvenile ontogeny (6 trials: 81 %, 5 trials: 10 %, 4 trials: 4 %, <4 trials: 5 %). From the first winter trials, we analysed the tests of the same individuals except for two individuals that had died and one male Satellite that had not been tested during the juvenile ontogeny (day 3 to 33 after hatching). From the second winter trials, we included all previously tested individuals that were still in our study population.

**Table 1:**
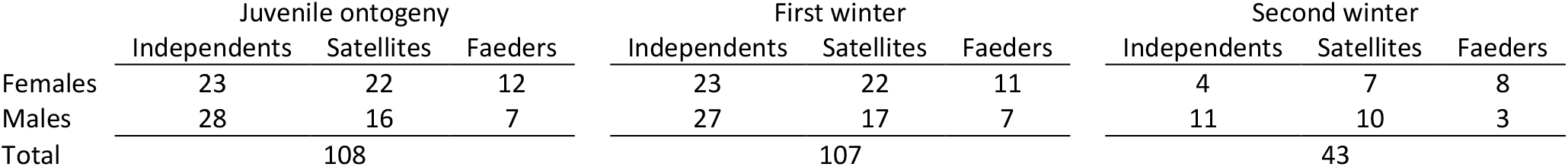
Sample sizes of each sex and morph for which the open field tests were analysed. Of the 109 individuals in total, one male Satellite was not tested during juvenile ontogeny. Individuals were tested up to six times during juvenile ontogeny (n = 607 trials), three times during the first winter (n = 321 trials), and three times during the second winter (n = 129 trials), resulting in a total of 1057 analysed open field tests.

We analysed the recorded videos of each trial using the Python-based automated tracking software TRex.run (Walter and Couzin 2021), which was run blind, i.e. without the information on bird ID, age, sex or morph information. The software provided the individual’s position (body centre) in each video frame (25 frames/sec). We used the adeHabitatLT package (version 0.3.27, Calenge 2006) in R (version 4.1.3, R Core Team 2022) to compute a smoothed trajectory of the birds walking on the floor, i.e., distance travelled. For smoothing, we subsampled the trajectory by taking only the coordinates of every 10^th^ frame and then reduced noise by removing movement distances of less than 1 cm between these subsampled frames (< 1 cm/0.4 sec). While being blind to the individuals’ IDs, we visually inspected the videos and manually excluded frames with tracking errors and periods where the bird flew higher than 0.5 m (flight periods). Individuals flew in 9.7 % of the videos, and the average duration of the time spent flying in these videos was 17.0 sec. Finally, we extracted the average travelling speed as the total distance travelled divided by the time spent on the ground. Based on the average travelling speed, we calculated the final response variable ‘distance travelled (in m) during a 10 min period’ as a proxy for activity, which was the response variable used in all subsequent analyses.

### Statistical Analyses

We conducted all statistical analyses in R (version 4.1.3, R Core Team 2022).

To analyse the development of activity during juvenile ontogeny (< 33 days of age), we implemented a Gamma Hurdle model (GHM) with cube-root transformed distance travelled as the response variable using the glmmTMB package (version 1.1.8, Brooks et al. 2017). GHMs consist of two parts, modelling two separate processes, and can be used to deal with zero-inflation, which we had in our juvenile ontogeny data. First, the hurdle part models the probability of not moving at all during the 10-minute tracking period (zero) vs. moving (one) as a binary process. Second, the conditional part models the non-zero values with a Gamma error distribution. For both parts we used the same fixed-effect structure, including sex (two-level factor: female, male), morph (three-level factor: Independent, Satellite, Faeder) and age (linear and quadratic continuous variable) centred on a mean of zero. We refrained from fitting a sex-morph interaction, because we did not have a clear prediction for it and fitting interactions can increase the risk for type I errors (Forstmeier and Schielzeth 2011; Gelman 2018). Further, we wanted to investigate the basal effects of sex and morph as the two main factors. The conditional part additionally included cohort (three-level factor: 2021, 2022, 2023) and ID (108 levels) as random intercepts. Finally, we included random slopes for ID over age to account for variation in individual developmental trajectories. We extracted the predicted values using the emmeans package (version 1.8.9, Lenth et al. 2023).

For the analysis of the mature phenotype of activity measured during first and second winter, we employed a Linear Mixed Model (LMM) with Gaussian error distribution using the lme4 package (version 1.1-35, Bates et al. 2015), since most birds moved during these trials and there was no zero inflation. Similar to the juvenile ontogeny analysis, we included sex, morph, winter (two-level factor: first, second) and trial number (three levels) as fixed effects, with cohort and ID (107 levels) as random effects.

To examine the developmental changes in the between-individual variation in activity, we calculated the adjusted individual repeatability (between-individual variance divided by the total phenotypic variance after accounting for fixed effects) using the rptR package (version 0.9.22, Stoffel et al. 2017). We classified the trials into four life stages, each consisting of three trials: (I) early (days 3-15) and (II) late (days 21-33) juvenile ontogeny, and (III) first and (IV) second winter (**Fehler! Verweisquelle konnte nicht gefunden werden**.). We calculated between individual variation as adjusted repeatability within each life stage as well as between life stages. Additionally, we investigated the “development” of repeatability during the juvenile ontogeny at a finer scale by calculating the repeatability for each combination of two juvenile ontogeny trials, similar to pairwise comparisons (Supplement, **Fehler! Verweisquelle konnte nicht gefunden werden**.). The formula for all underlying Gaussian LMMs and adjusted repeatabilities included cube-root transformed distance travelled as the response variable, sex, morph and trial as a fixed and cohort and ID as random effects. Furthermore, we used the variance partitioned between ID and the residual variance in the four life-stage LMMs as indicators of changes in between- and within-individual variation, respectively.

We conducted two supplemental analyses regarding the development of behavioural stability. First, we investigated whether individuals establish a stable ranking within sex-morph groups based on the distance travelled throughout the juvenile ontogeny. This allowed us to assess the consistency of individual phenotypes for a trait that changes during juvenile development. High rank stability indicated that e.g., more active chicks were consistently more active relative to their peers over consecutive trials (Supplement, **Fehler! Verweisquelle konnte nicht gefunden werden**.). Second, we examined from which age on the chick behaviour can predict the mature phenotype of activity measured in the first winter (Supplement, **Fehler! Verweisquelle konnte nicht gefunden werden**.). Both analyses confirmed the results of the repeatability analysis.

## Results

### Development of activity

Distance travelled showed a clear relationship with the age of young ruffs during the juvenile ontogeny period. Initially on day 3, the birds travelled a median distance of 0.23 m (range: 0 – 30.13 m) within the 10 min test duration. Median distance travelled increased to 19.07 m (range: 0 – 257.93 m) on day 33. Notably, the increase in distance travelled over age was not linear but showed a better fit to a quadratic trajectory. The distance travelled initially decreased from day 3 to day 9 but then showed a strong increase from day 21 onwards (Table 2, Figure 1 A, **Fehler! Verweisquelle konnte nicht gefunden werden**.). Similarly, the probability of zero movement followed a quadratic relationship with a slight initial increase followed by a decrease towards zero (Table 2, **Fehler! Verweisquelle konnte nicht gefunden werden**. B). The young ruffs were more likely to move than to stay immobile from day 21 onwards (probability of zero movement < 0.5).

**Table 2:**
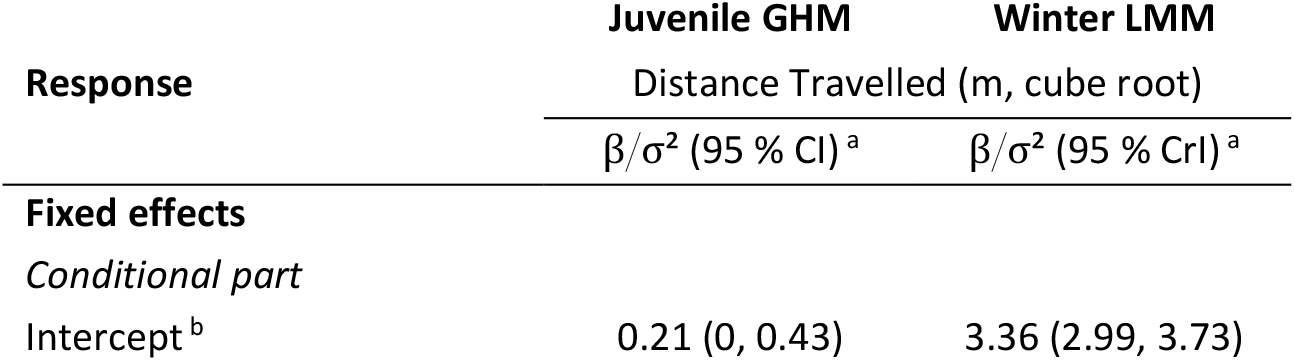
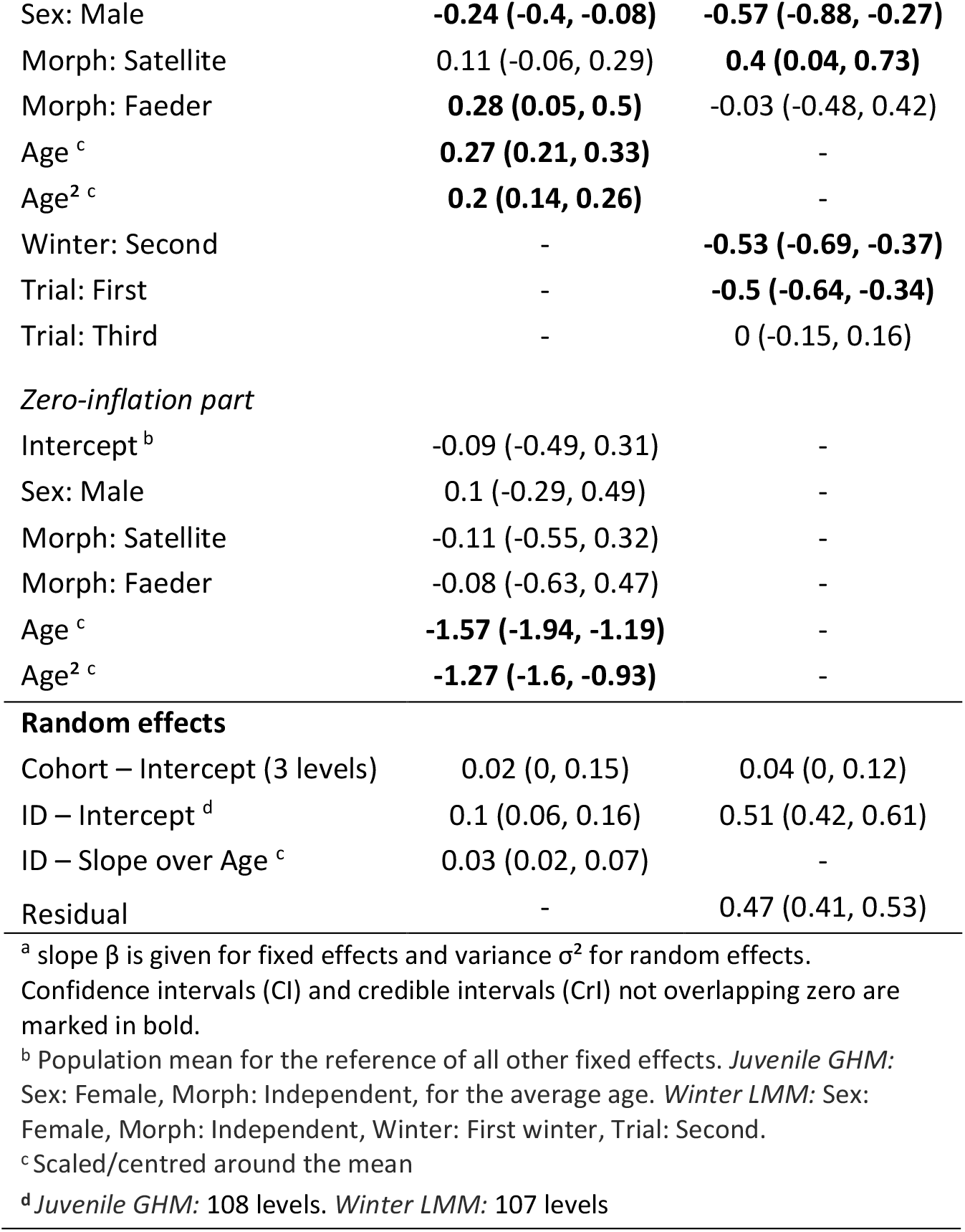
Predictors of activity in young ruffs during the juvenile ontogeny period (days 3-33) and during the winter (first and second winter). Shown are the outputs of the Juvenile Gamma Hurdle Model (GHM) based on distances travelled of 108 individuals (607 trials) and of the Winter Linear Mixed Model (LMM) on 107 individuals (450 trials), respectively.

| Response | Juvenile GHM | Winter LMM |
| --- | --- | --- |
|  | Distance Travelled (m, cube root) |  |
| | $\beta/\sigma^2$ (95 % CI)<sup>a</sup> | $\beta/\sigma^2$ (95 % CrI)<sup>a</sup> |
| **Fixed effects** |  |  |
| *Conditional part* |  |  |
| Intercept<sup>b</sup> | 0.21 (0, 0.43) | 3.36 (2.99, 3.73) |
| Sex: Male | **-0.24 (-0.4, -0.08)** | **-0.57 (-0.88, -0.27)** |
| Morph: Satellite | 0.11 (-0.06, 0.29) | **0.4 (0.04, 0.73)** |
| Morph: Faeder | **0.28 (0.05, 0.5)** | -0.03 (-0.48, 0.42) |
| Age<sup>c</sup> | **0.27 (0.21, 0.33)** | - |
| Age<sup>2 c</sup> | **0.2 (0.14, 0.26)** | - |
| Winter: Second | - | **-0.53 (-0.69, -0.37)** |
| Trial: First | - | **-0.5 (-0.64, -0.34)** |
| Trial: Third | - | 0 (-0.15, 0.16) |
| Zero-inflation part |  |  |
| Intercept<sup>b</sup> | -0.09 (-0.49, 0.31) | - |
| Sex: Male | 0.1 (-0.29, 0.49) | - |
| Morph: Satellite | -0.11 (-0.55, 0.32) | - |
| Morph: Faeder | -0.08 (-0.63, 0.47) | - |
| Age<sup>c</sup> | **-1.57 (-1.94, -1.19)** | - |
| Age<sup>2 c</sup> | **-1.27 (-1.6, -0.93)** | - |
| Random effects |  |  |
| Cohort – Intercept (3 levels) | 0.02 (0, 0.15) | 0.04 (0, 0.12) |
| ID – Intercept<sup>d</sup> | 0.1 (0.06, 0.16) | 0.51 (0.42, 0.61) |
| ID – Slope over Age<sup>c</sup> | 0.03 (0.02, 0.07) | - |
| Residual | - | 0.47 (0.41, 0.53) |

<sup>a</sup> slope $\beta$ is given for fixed effects and variance $\sigma^2$ for random effects.
Confidence intervals (CI) and credible intervals (CrI) not overlapping zero are marked in bold.
<sup>b</sup> Population mean for the reference of all other fixed effects. *Juvenile GHM*:
Sex: Female, Morph: Independent, for the average age. *Winter LMM*: Sex:
Female, Morph: Independent, Winter: First winter, Trial: Second.
<sup>c</sup> Scaled/centred around the mean
<sup>d</sup> *Juvenile GHM*: 108 levels. *Winter LMM*: 107 levels

**Figure 1:**
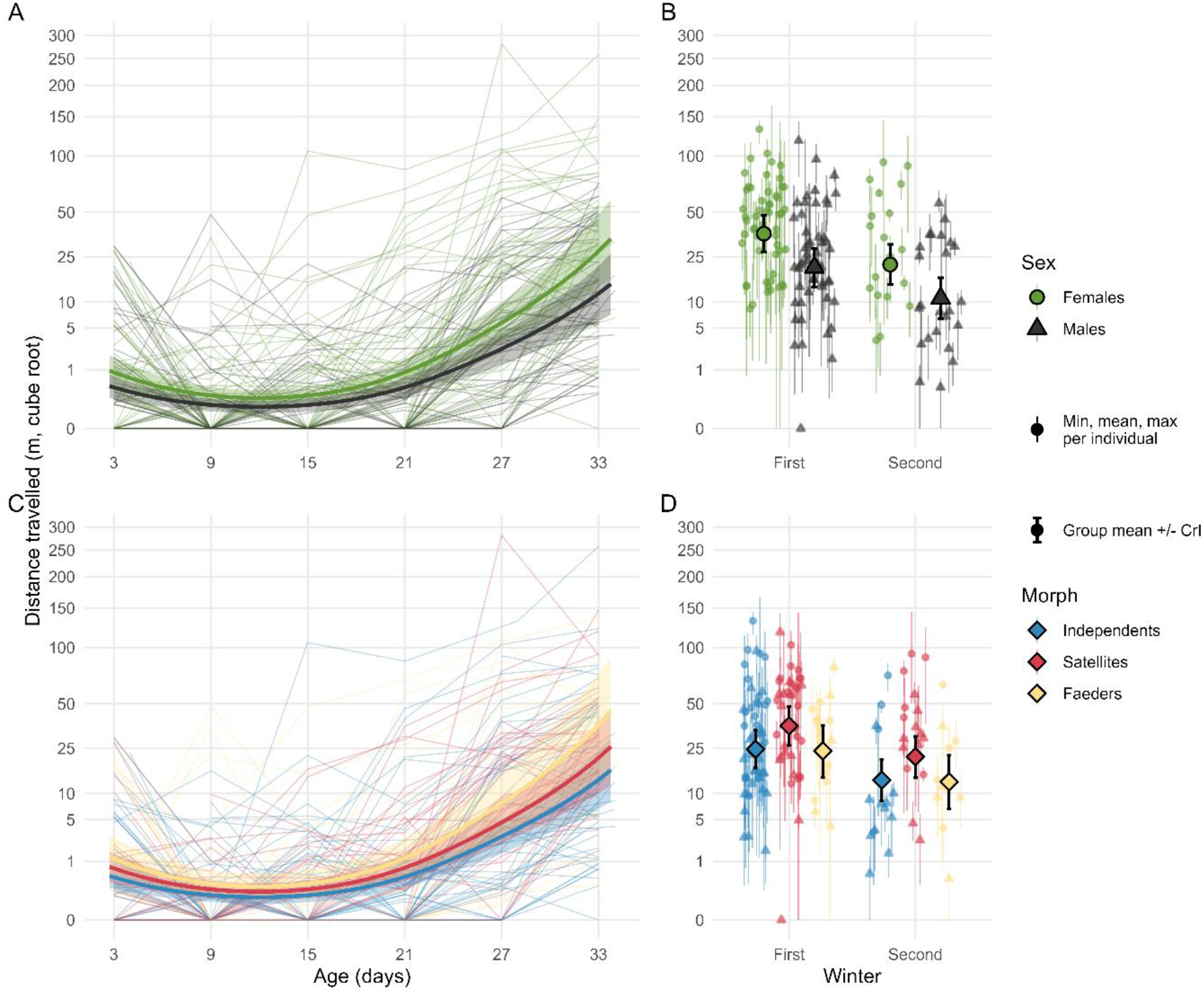
Sex and morph differences in activity in young ruffs during different life stages. A&C) Juvenile Hurdle Gamma Model: Development of distance travelled during juvenile ontogeny (day 3 until day 33). Curves indicate A) sex differences (statistically clear) and C) morph differences (clear difference only between Independents and Faeders). In A&C, the predicted curves are plotted as thick lines with 95 % confidence intervals (shaded areas). The raw data are plotted as thinner lines in the background. B&D) Winter Linear Mixed Model: Mature phenotype of activity measured during the birds’ first and second winter. B) Sex differences in first and second winter (clear differences between sexes and winters). D) Morph variation in first and second winter (clear difference between Independents and Satellites). In B&D, predicted group means are plotted as solid shapes outlined in black, with error bars indicating the 95 % credible intervals. The raw data are plotted as semi-transparent point ranges indicating the minimum, mean and maximum values measured for each individual. Colours indicate either the two sexes (females in green and males in black), or the three morphs (Independents in blue, Satellites in red and Faeders in yellow). Shapes indicate the two sexes (females as circles and males as triangles) and the morphs (diamonds, same for all three morphs).

In their first winter, the birds covered a median distance of 30.19 m (range: 0 – 166.54 m) during the 10 min test duration. Thus, the activity levels achieved at the end of the juvenile ontogeny already covered a similar range of values as the ones measured during the first winter. In comparison, the individuals moved less during their second winter (mean predicted difference of 12.13 m; Table 2, Figure 1 B). At this age, the birds travelled a median distance of 15.03 m (range: 0 – 144.96 m).

### Sex and morph differences in activity

Comparing the sexes, males generally travelled shorter distances than females during early juvenile ontogeny (Table 2, Figure 1 A). On day 33, the predicted sex difference in travel distance was the largest with 14.48 m (smallest difference on day 9 = 0.26 m). However, the likelihood of moving vs. not-moving was equal for both sexes (Table 2, **Fehler! Verweisquelle konnte nicht gefunden werden**. B). Additionally, males generally travelled shorter distances than females during the first and second winter (predicted sex difference of 13.26 m, averaged over both winters; Table 2, Figure 1 B).

Differences between morphs were generally not as clear and not as consistent as those between the sexes. Throughout the juvenile ontogeny, Satellites exhibited no clear differences in activity compared to Independents. However, Faeders travelled notably longer distances than both other morphs (largest predicted difference on day 33 = 24.65 m, smallest difference on day 9 = 0.46 m; Table 2, Figure 1 C). The probability of zero movement did not vary between the three morphs (Table 2, **Fehler! Verweisquelle konnte nicht gefunden werden**. B). In the first and second winter, Satellites moved further than the other two morphs (predicted difference of 9.69 m, averaged over both sexes) whereas Independents and Faeders travelled similar distances (Table 2, Figure 1 D).

### Development of between-individual differences in activity

Adjusted repeatability increased across the four life stages from R = 0.11 during early juvenile ontogeny to R = 0.54 in the first and R = 0.48 in the second winter (Figure 2, on the diagonal), indicating high repeatability of the measured behaviour in the mature phenotype (Bell et al. 2009; Holtmann et al. 2017; Dingemanse and Wright 2020). The same general pattern of increasing repeatability with age was evident comparing repeatabilities between life stages (Figure 2, below the diagonal). The variance partitioning in the corresponding LMMs indicated the general pattern that the increase in repeatability was driven by an increase in between-individual and a decrease in within-individual variation with age (**Fehler! Verweisquelle konnte nicht gefunden werden**., **Fehler! Verweisquelle konnte nicht gefunden werden**.). Similarly, rank stability increased throughout juvenile ontogeny (**Fehler! Verweisquelle konnte nicht gefunden werden**., **Fehler! Verweisquelle konnte nicht gefunden werden**.) and from day 9 onwards, chick activity predicted the mature phenotype, i.e., the activity observed in their first winter, increasingly well. The measurements on days 27 and 33 emerged as the best predictors of the mature phenotype (**Fehler! Verweisquelle konnte nicht gefunden werden**., **Fehler! Verweisquelle konnte nicht gefunden werden**.).

**Figure 2:**
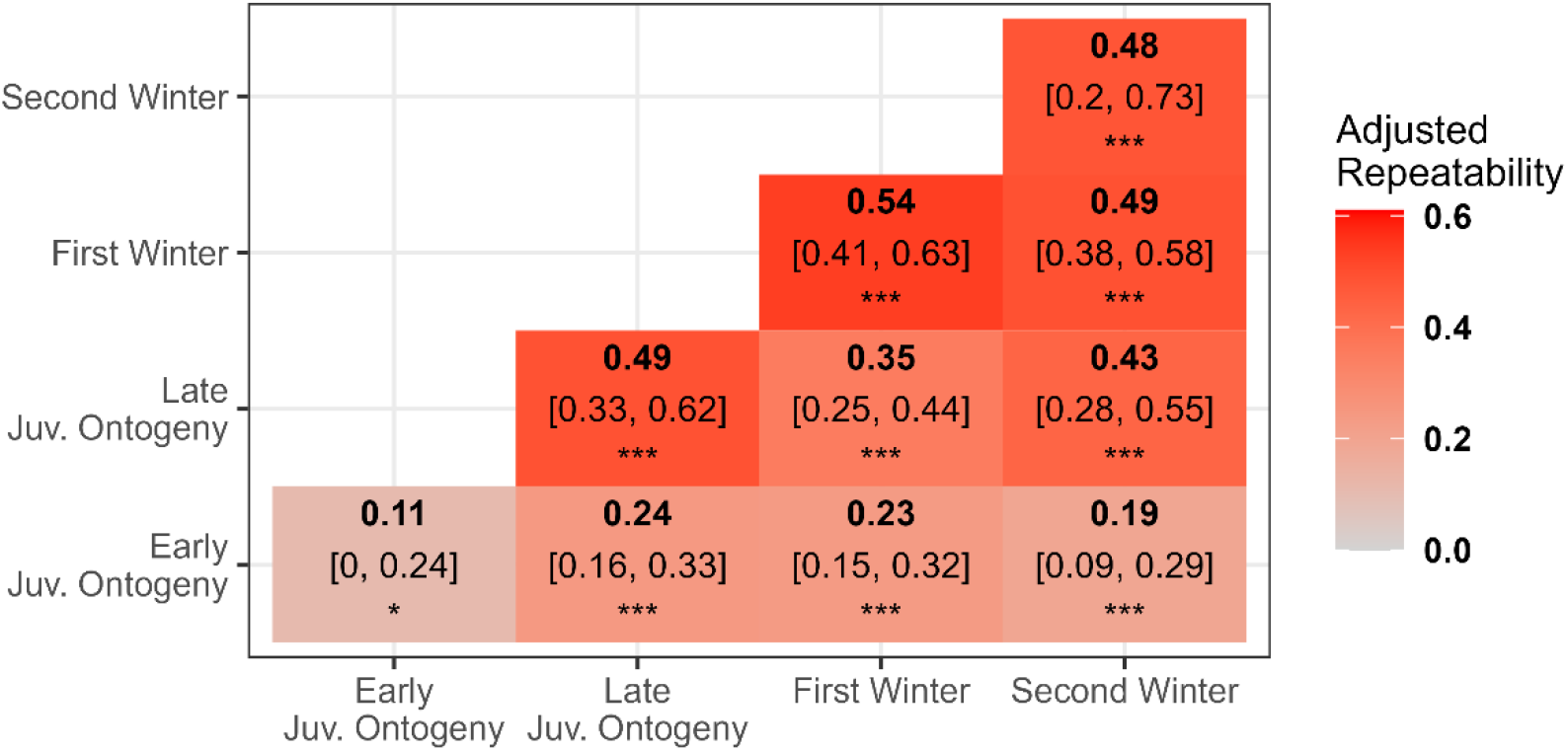
Adjusted repeatabilities of activity within and between all life stages tested. The diagonal contains repeatabilities of distances travelled within life stages of three trials each. The lower part of the matrix contains repeatabilities of all pairwise combinations between life stages. Adjusted repeatabilities are shown in bold, [95 % confidence interval] and significance is given as p <0.001 ***, and p < 0.05 *.

## Discussion

Intraspecific variation plays a major role for the adaptive potential of populations. However, the extent of intraspecific variation during juvenile ontogeny and developmental patterns are often unknown. We examined intraspecific behavioural variation in the development of activity in an unfamiliar environment, a movement-related behavioural trait, in precocial ruffs. We investigated how sex, morph and individual variation shape activity of young ruffs. Our study had three main findings. First, as predicted, activity generally increased with age during the early ontogeny when the young ruffs were growing fast. During the first month, activity rapidly increased after the developing chicks shed their crouching reflex in response to a potential threat. After one month, when their physical growth was largely complete, the birds reached the activity level that remained largely stable in their first and second winter. Second, differences in activity were greater between sexes than between morphs. Overall, females travelled longer distances than males. In contrast, we did not detect consistent morph differences that remained stable over time. Third, as predicted, individual repeatability increased during early ontogeny. Repeatability was initially low, but increased throughout juvenile ontogeny and remained high (∼0.5) within and between first and second winter measurements.

### Development of activity

Throughout the first month of life, we observed a non-linear increase in the activity of young ruffs. Despite their physical ability to locomote, the chicks showed low levels of activity during the first three trials, until day 15, and typically remained motionless on the ground in a crouching position. This behaviour is a common response of young precocial chicks to potential threats, where they rely on their cryptic plumage to evade potential predators (Simmons 1955; Colwell 2010; Volkmer et al. 2024). Interestingly, the crouching time increased between first and second trial (ages 3 and 9 days). During the first trial on day 3, the chicks were still more likely to move, resulting in a slightly higher average activity compared to day 9 or 15. During this first trial, several chicks started to walk while calling loudly soon after the start of the trial (personal observation). One explanation for this is that the long trial duration (10 mins) induced calling in the isolated chicks to establish contact with their family members. Chicks at this age are not yet self-sufficient in thermoregulation and require regular brooding (Visser and Ricklefs 1993; Colwell 2010). By day 9, thermoregulation appeared to be less of an issue and chicks were more likely to remain motionless throughout the 10-minute trial resulting in the overall lowest activity levels observed.

As physical growth continued and mobility increased, older chicks shed the crouching reflex leading to a notable increase in activity in the second half of the juvenile ontogeny period. The most substantial increase in average activity occurred from day 27 onwards. By this age, nearly all chicks had already fledged. With *ad libitum* food supply the chicks typically gain flight ability between day 16 and day 25 (Giraldo-Deck 2022). By day 33, activity levels of young ruffs reached those levels observed in the first winter. In comparison with the first winter, we noted a decrease in activity during the second winter. One explanation for the reduced activity might be increased habituation to regular handling in captivity. Handling by humans has been shown to reduce activity in open field tests in birds (Feenders et al. 2011). Over the years, the ruffs may become more relaxed and consequently less affected by the unfamiliar situation to which they are exposed during the open field test, resulting in lower activity levels.

### Sex differences in activity

Throughout juvenile ontogeny, females consistently travelled longer distances than males and this difference persisted throughout the winter tests. While the crouching reflex was a predominant effect during the first half of this period and the likelihood of moving vs. not-moving was equal for both sexes, the sex difference became especially apparent during the second half of juvenile ontogeny, once the majority of the chicks started to move. The observed sex difference in activity is consistent with findings in other species such as rats and sheep, where females have been shown to travel longer distances in open field tests (e.g. Knight et al. 2021; Blizard et al. 1975; Vandenheede and Bouissou 1993). However, a recent meta-analysis did not detect a general sex difference in the personality trait category ‘exploration-avoidance’ (as defined by Réale et al. 2007), either across various animal taxa, or across a wide range of bird species (Harrison et al. 2022). Further, this study did not corroborate the hypothesis that sexual size dimorphism would affect exploration behaviour (Harrison et al. 2022). However, the meta-analysis did not specifically evaluate differences during development. During this time, sexual size dimorphism may show a stronger relation to such activity-related behaviours. In ruffs, the size differences between males and females emerge already during the first week of life (Giraldo-Deck et al. 2020). These sex differences may have consequences for activity. First, although males have higher absolute growth rates than females, females have higher relative growth rates meaning that they approach their final body size faster than males (Giraldo-Deck et al. 2020). The faster relative growth may lead to a faster development of mobility in females than in males (Giraldo-Deck et al. 2020; Giraldo-Deck 2022). Second, especially males, the larger sex, might face a trade-off between allocating energy reserves to growth versus activity. The larger sex is typically more vulnerable to adverse environmental conditions, such as limited food availability, during early ontogeny (Clutton-Brock et al. 1985; Blanckenhorn 2005; Loonstra et al. 2018). With a significant proportion of their resources allocated to growth (Clutton-Brock et al. 1985), males should hence likely have fewer energy reserves available for movement (Careau et al. 2008). Consequently, males may expend less energy on locomotion and spend more time resting to conserve energy. A similar energy allocation trade-off may also underlie the sex differences in activity that persist into the first and second winter. Even after growth is complete, males seemed to show greater inertia, seemingly prioritizing energy conservation. Males may be less inclined to move and more selective about when to initiate movement, even in potentially threatening situations, like our activity test situation. Alternatively, the higher activity observed in ruff females may also be related to known sex differences in migration distance. Female ruffs overwinter predominantly in Africa and South Asia whereas males typically stay further north (Delaney et al. 2009; Jaatinen et al. 2010). This phenomenon is consistent with the known increase in flight costs with body mass (Hein et al. 2012), and similar patterns have been observed in other size-dimorphic species, where the larger sex tends to exhibit lower levels of general activity (Ruckstuhl 1998; Ruckstuhl and Neuhaus 2000).

### Morph differences in activity

The variation in activity between the three morphs was not as clear as between the sexes. Faeders appeared to display somewhat higher activity levels during juvenile ontogeny compared to Independents and Satellites (Table 2). **Fehler! Verweisquelle konnte nicht gefunden werden**.Further, Faeders did not display higher activity levels during the first and second winter. In line with the smaller females moving more, also the higher activity of Faeders during juvenile ontogeny might be related to their smaller body size compared to the other two morphs. However, these differences were not found during the winter trials and therefore we refrain from further interpretation. At the mature age, Satellites exhibited notably higher levels of activity than the other two morphs. However, we note that alternative models, i.e., a LMM on the first winter data alone did not show these differences (**Fehler! Verweisquelle konnte nicht gefunden werden**.). The higher activity of Satellites in the second winter data may not hold due to the substantially lower sample size (**Fehler! Verweisquelle konnte nicht gefunden werden**.). Therefore, it remains unclear whether this pattern is robust. Given that the genetic differences between morphs are attributed to a relatively small autosomal inversion comprising only approximately 100 genes (Küpper et al. 2016; Lamichhaney et al. 2016), activity may not be affected by the genetic differences between morphs.

Comparisons with other species that exhibit ARTs reveal no clear pattern regarding the relationship between activity in unfamiliar environments and mating morphs. As in our study, in some species, activity appears to be unrelated to mating morphs (Wilson and Kelly 2019; Rochais et al. 2021; Warrington et al. 2022). Conversely, in other systems, ARTs are associated with differences in activity (e.g. Scantlebury et al. 2008; Han and Jablonski 2019; Madrid et al. 2020) or other personality traits (e.g. Barcelo-Serra et al. 2020; Rochais et al. 2021; Brock et al. 2022; Holtmann et al. 2022). We conclude from this that evidence for behavioural differences outside the mating context in species with ARTs remains limited.

### Development of between-individual differences in activity

The repeatability of activity increased with age during juvenile ontogeny stages. Similarly, the individuals became more stable in their ranking within the sex-morph groups (Supplement, **Fehler! Verweisquelle konnte nicht gefunden werden**.), and their activity during juvenile ontogeny predicted the mature phenotype increasingly well (Supplement, **Fehler! Verweisquelle konnte nicht gefunden werden**.). By the first winter, repeatability reached high levels, indicating increasing stability of the behavioural phenotypes in this personality trait. This increase in repeatability seems to be attributed to both a decrease in within-individual variation and an increase in between-individual variation, a phenomenon known as fanning-out between individual trajectories (Stamps and Groothuis 2010; Sih et al. 2015; Fisher et al. 2018). Generally, during juvenile ontogeny, age-related physical growth and development of mobility emerge as the primary determinants of variation in activity. As individuals mature, the influence of age decreases and personality-related individual variation becomes more pronounced.

Our findings together with other empirical studies (e.g. Freund et al. 2013; Fisher et al. 2015; Polverino et al. 2016; Carlson and Tetzlaff 2020) are consistent with theoretical predictions that individuals diverge in their behavioural tendencies as they develop and eventually become more consistent in their behaviour (Duckworth 2010; Stamps and Krishnan 2017). This developmental trend, characterised by higher within-individual variation in younger individuals, has been attributed to inexperience (Polverino et al. 2016; Ersoy et al. 2024). Our results are consistent with previous hypotheses that individual exploration behaviour, analogous to the activity measured in this study, partially develops during early juvenile ontogeny, with further refinement occurring from juvenile to adult stages (Kok et al. 2019; Ersoy et al. 2024).

Despite the controlled environment in which our birds were raised and kept, a wide range of variation in activity emerged. It is not clear how the experimental set up affected between and within individual variation. On the one hand, variation in our study may be lower than in the wild due to reduced stimulation and environmental influences (Groothuis and Trillmich 2011; Laskowski et al. 2022). On the other hand, variation in activity in our study may be larger than in the wild due to the absence of time or environmental constraints and/or predation pressure, as relaxed predation pressure has been shown to lead to increased behavioural variability in prey species (Dingemanse et al. 2009; Laskowski and Bell 2013). In the latter case, motivation to move has been shown to vary between individuals (Hertel et al. 2023). However, the activity trait we measured may not solely reflect intrinsic, self-motivated movement behaviour. The unfamiliar test situation might be perceived as threatening and elicit an externally induced stress, fear, or anxiety-related response, causing similar individual variation along the freeze vs. flight or proactive vs. reactive continuum (Carter et al. 2013; Perals et al. 2017). This ambiguity regarding the interpretability of the activity trait as an externally induced stress response versus an internally motivated exploratory behaviour has raised increasing concerns within the field of animal personality research, where the behaviour has traditionally been termed ‘exploration behaviour’. Furthermore, it is questionable whether the measured behaviour really refers to an individual’s response to a novel environment as many studies, including ours, reuse the same test arena for subsequent tests of the same individual, leading to a decline in novelty with repeated exposure (e.g., Dingemanse et al. 2002; Dingemanse et al. 2012; Bijleveld et al. 2014; Wuerz and Krüger 2015; Minderman et al. 2010). Therefore, we use ‘activity’ as a purely descriptive term in the context of an unfamiliar environment.

## Conclusion

In conclusion, our experimental assessment of the variation in activity measured in open field tests indicates that this trait may represent a personality trait in ruffs that develops and stabilizes during the first month after hatching and then remains repeatable over years. By the end of juvenile ontogeny, the birds already reach activity levels observed in the first winter and their activity already predicts the long-term stable, mature phenotype.

Activity was clearly related to sex but not to morph. Overall, females were significantly more active in the unfamiliar environment, which may be explained by the strong sexual size dimorphism and energetic constraints and trade-offs that limit male activity. Whether activity differs between the morphs remains less clear. The lack of consistent differences suggests that the supergene that underlies the three mating morphs does not have an overly strong effect on activity-related traits. However, it is too early to conclude that supergenes underlying mating behaviours do not impact personality traits that are seemingly unrelated with reproduction and we call for further studies to investigate how sexes and morphs affect personality traits.

## Acknowledgements

We thank Claudia Scheicher and Petra Neubauer for help with animal care and Melanie Schneider for genetic sex and morph determination. We also thank Nikolas Heinecke for technical help with the camera equipment. Special thanks also to Zsombor Fueloep for assisting with python challenges in the automated tracking process. We are grateful to Niels Dingemanse and members of the research group Behavioural Genetics and Evolutionary Ecology for discussion of the results and constructive feedback.

## Ethics approval statement

All procedures involving animals were conducted in accordance with the ethical guidelines of the [Institutional Animal Care and Use Committee (IACUC) or equivalent]. All necessary permits for the breeding and housing of the birds were obtained. No animals were harmed during the experiments.

## Data availability

Ten exemplary raw videos (to not exceed storage capacity), all data tables containing the coordinates of the raw trajectories, a table containing the birds’ information and scripts are stored in Edmond the Open Research Data Repository of the Max Planck Society (https ://).

## Author contributions (CRediT)

**Veronika A. Rohr-Bender**: conceptualization (equal), data curation (lead), formal analysis (lead), funding acquisition (equal), investigation (lead), methodology (lead), visualization (lead), writing – original draft preparation, review & editing (lead). **Krisztina Kupán**: formal analysis (support), methodology (support), writing – review & editing (support). **Guadalupe Lopez-Nava**: data curation (support), formal analysis (support), methodology (support), writing – review & editing (support). **Wolfgang Forstmeier**: formal analysis (support), methodology (support), writing – review & editing (support). **Anne Hertel**: formal analysis (support), methodology (support), writing – review & editing (support). **Vitali Razumov**: investigation (support), writing – review & editing (support). **Katrin Martin**: investigation (support), writing – review & editing (support). **Bart Kempenaers**: resources (support), writing – review & editing (support). **Clemens Küpper**: conceptualization (equal), formal analysis (support), funding acquisition (equal), methodology (support), resources (lead), supervision (lead), writing – original draft preparation (support), review & editing (support).

