## Supplement for "Sex and morph variation in activity from early ontogeny to maturity in ruffs (*Calidris pugnax*)"

### Supplementary material

#### Individuals tested for the experiment

To investigate the development of activity in an unfamiliar environment early in life, we conducted a series of open field tests with young ruffs that were raised in captivity over three consecutive breeding seasons (2021-2023). We tested the birds from day 3 after hatching until day 33, when all juveniles had fledged. In addition, we tested the ruffs again during their first and second winter to examine the behavioural stability of their mature phenotype. For 3% of the scheduled trials, we shifted individual tests by one day. Chicks that appeared unwell on a test day were not tested meaning that 24% of the 271 test individuals who survived until day 33 (end of juvenile ontogeny) did not undergo all six open field tests (Figure S1). We chose a subset of 109 individuals to be analysed further, as we aimed for a balanced representation of all sex-morph classes (Figure S1, **Fehler! Verweisquelle konnte nicht gefunden werden.**).

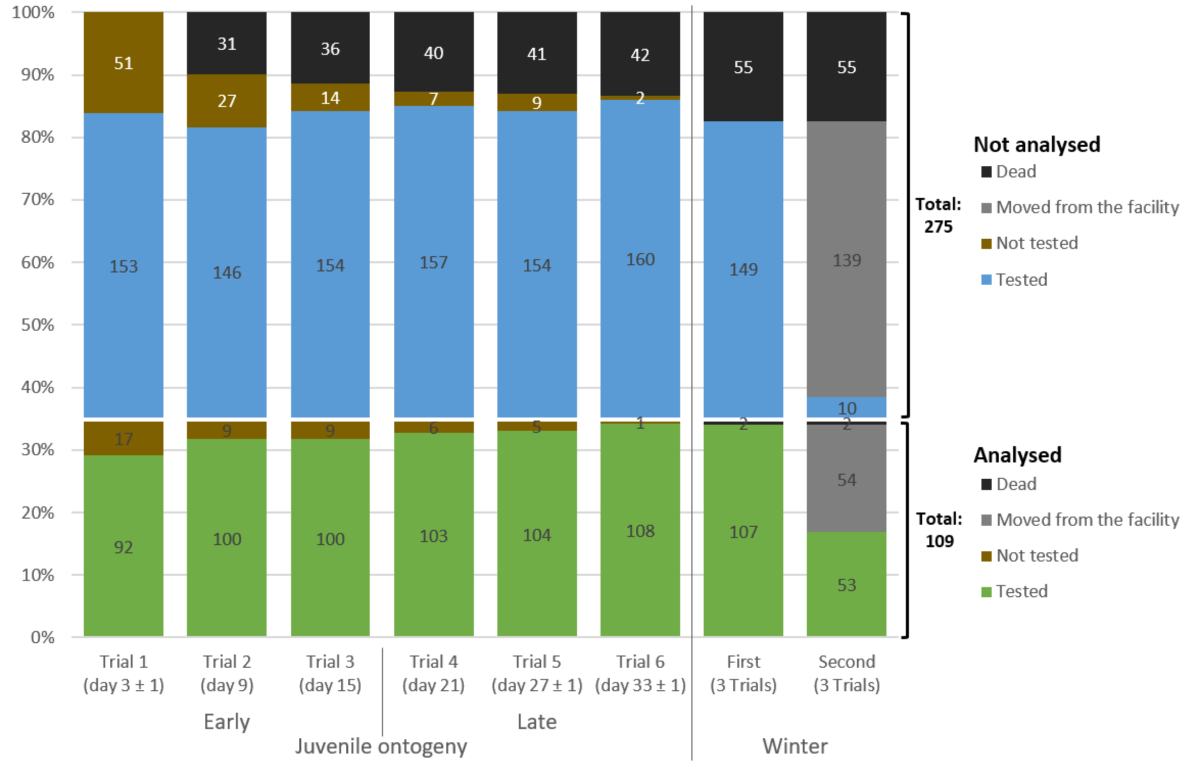


*Figure S1: Sample sizes of individuals that were tested and analysed during the measured life stages. In total, 292 trials of 103 individuals were analysed for the life stage of early juvenile ontogeny, 315 trials of 108 individuals for late juvenile ontogeny, 321 trials of 107 individuals for the first winter and 129 trials of 43 individuals for the second winter.*

### Juvenile GHM

As the young chicks did not move in many of the trials, which caused a zero-inflation, we measured during juvenile ontogeny we analysed this life stage with a Gamma Hurdle Model (GHM). It consists of a conditional part focusing on the data points being non-zero (Figure S2 A, Figure S3 A). Separately, it treats the zero-inflation in the data with a binomial model considering zero vs. non-zero values (Figure S2 B, Figure S3 B). See main manuscript for more detailed description and discussion.

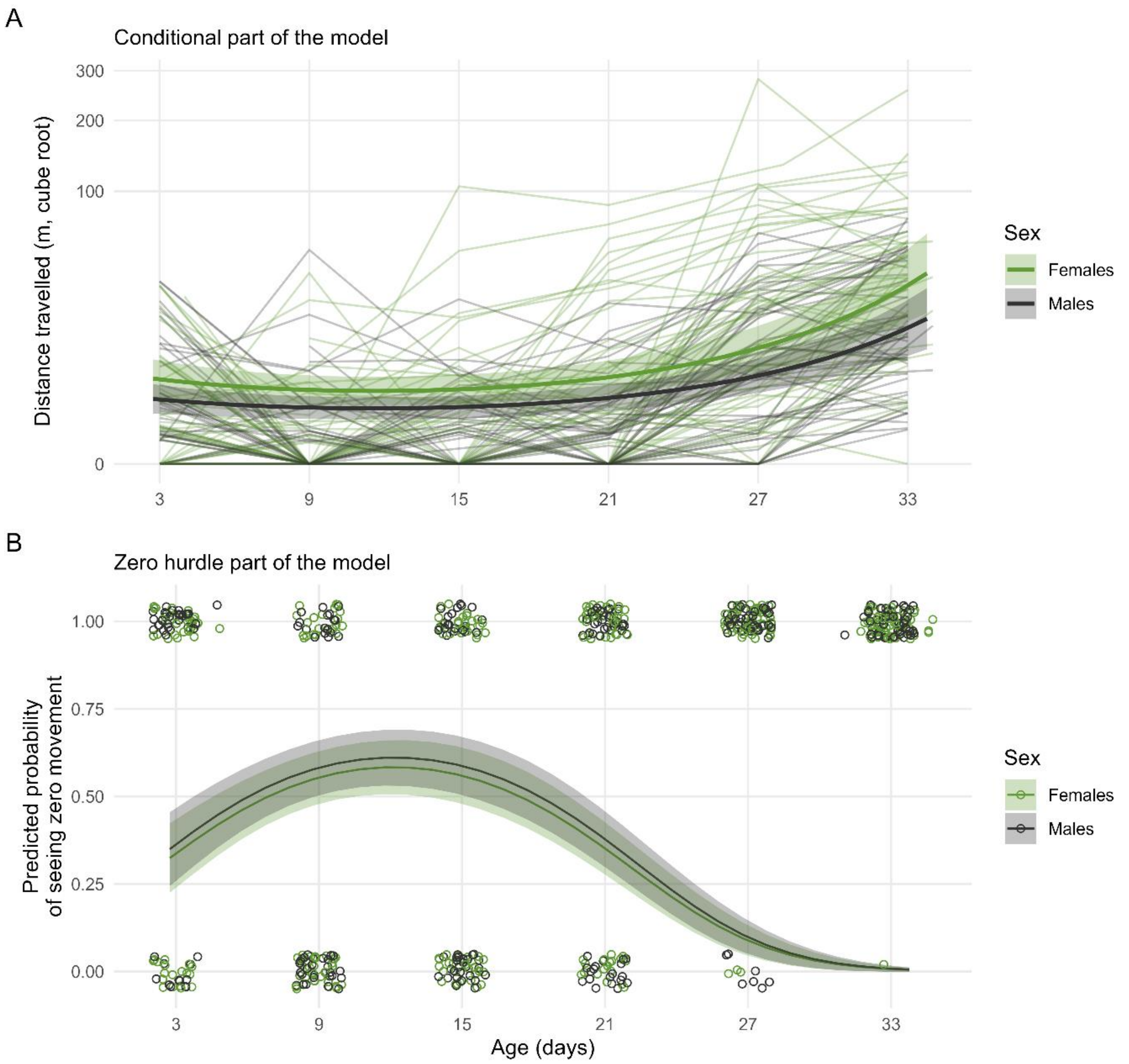


*Figure S2: Juvenile GHM: Sex differences in the ontogeny of activity (distance travelled) from day 3 until day 33. A) The predicted curves of the conditional part of the model are plotted as thick lines with 95 % confidence intervals (shaded areas) for females in green and males in black. The raw data is plotted as the thinner lines in the background, each connecting all measurements of one individual. The y-axis is displayed with cube root transformation. B) The predicted probability of seeing zero movement is displayed as thick lines with 95 % confidence intervals. The raw data points are displayed as being zero vs. non-zero (value = 1.00). Jitter was added to increase visibility of data points.*

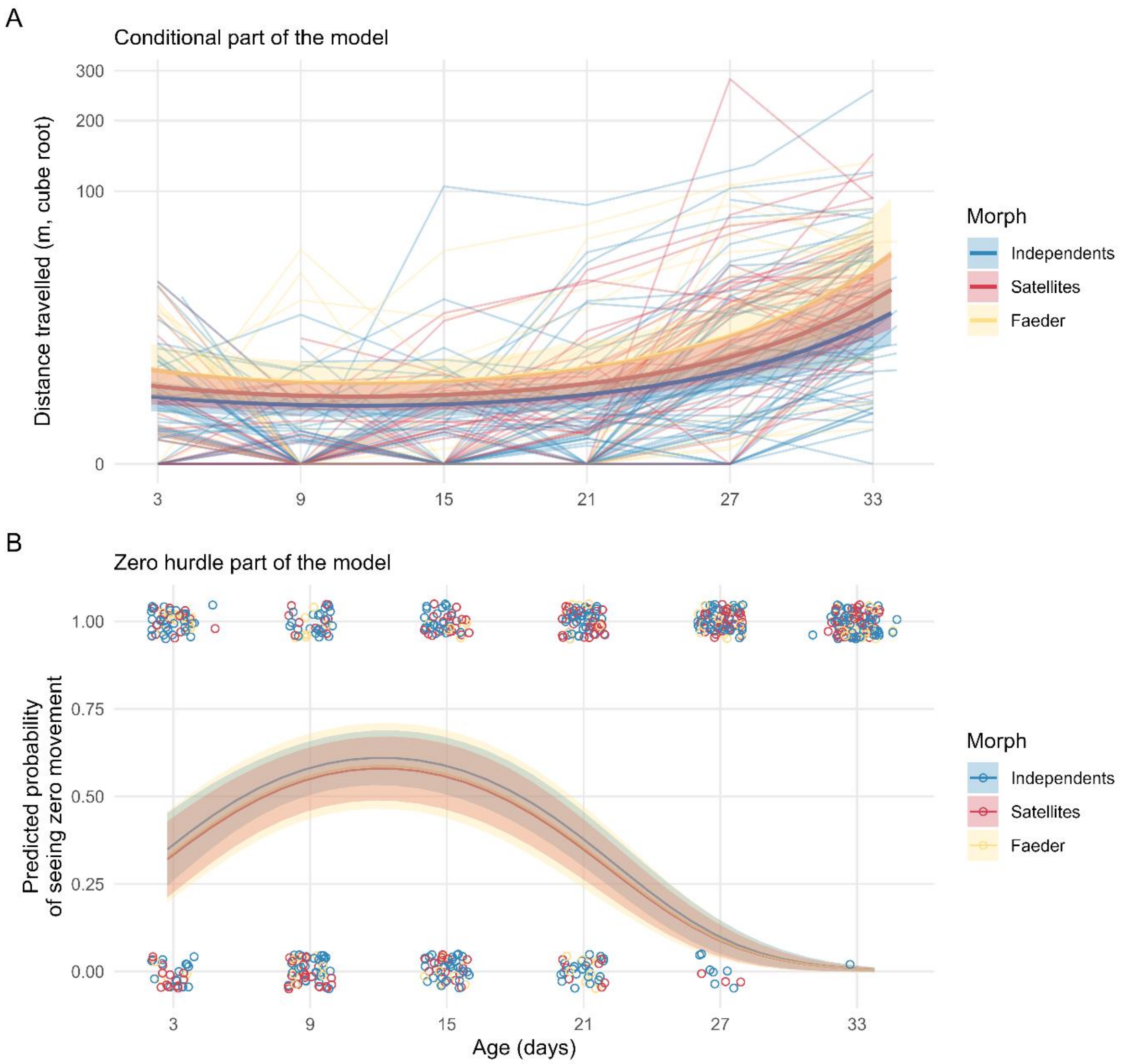


*Figure S3: Juvenile GHM: Sex and morph differences in the ontogeny of activity (distance travelled) from day 3 until day 33. A) The predicted curves of the conditional part of the model are plotted as thick lines with 95 % confidence intervals (shaded areas) for Independents in blue, Satellites in red and Faeders in yellow. Females and males are shown in separate panels. The raw data is plotted as the thinner lines in the background, each connecting all measurements of one individual. The y-axis is displayed with cube root transformation. B) The predicted probability of seeing zero movement is displayed as thick lines with 95 % confidence intervals. The raw data points are displayed as being zero vs. non-zero (value = 1.00). Jitter was added to increase visibility of data points.*

**Life stage LMMs**

We classified all trials measured into four life stages, each consisting of three trials: (I) early and (II) late juvenile ontogeny, (III) first and (IV) second winter (Figure S1). We ran a LMM separate for each life stage (Life stage LMMs) and used the variance partitioned between ID and the residual variance as indicators of changes between life stages in between- and within-individual variation, respectively (Table S1).

The strong age effect reflected by the Juvenile GHM and LMM is also visible in the LMMs on early and late juvenile ontogeny. The early juvenile ontogeny LMM reports higher activity in the first trial (age 3 days) compared to trial two and three (age 9 and 15 days). The chicks being more active in the first trial can again be interpreted as them seeking contact to their family as they still lack the ability to thermoregulate independently due to their young age. The low activity at trial two and three again indicates the strong crouching down reflex. The late juvenile ontogeny LMM also reports a steady increase in activity from the first trial (age 21 days) to trial three (age 33 days; Table S1).

Comparing the variance partitioning in the four life stages (Table S2), reveals a general increase of variance explained by ID, which we interpret as a general increase of between-individual variation. Accordingly, the variance explained by the residuals generally decreases over the four life stages, which indicates a decrease of within-individual variation. Cohort usually only explains a minor part of the total variance. The slightly higher percentage of variance explained by the cohort during late juvenile ontogeny might be due to a generally lower amount of variance being left unexplained by the fixed effects in that life stage. The high percentage explained by it in the second winter likely occurs due to the small data set, which additionally is unbalanced regarding cohort, sex and morph. We are aware that the values might not be directly comparable, because the variance explained by the fixed effects might vary between life stages, and especially in the second winter the sample size is substantially lower (Figure S1), however, the general pattern holds.

See main manuscript for further description and discussion.

*Table S1: Output of four Linear Mixed Models (LMM) considering the four measured life stages separately.*

| | **LMM – early juv.-ontogeny** | **LMM – late Juv.-ontogeny** | **LMM – first winter** | **LMM – second winter** |
|---|---|---|---|---|
| **Response** | Distance travelled (m, cube root) | | | |
| | $\beta/\sigma^2$ (95 % CrI) [a] | | | |
| **Fixed effects** | | | | |
| Intercept [b] | 0.46 (0.2, 0.71) | 2.1 (1.5, 2.69) | 3.41 (3.06, 3.76) | 2.58 (1.62, 3.48) |
| Sex: Male | -0.07 (-0.29, 0.15) | **-0.62 (-1.02, -0.26)** | **-0.55 (-0.88, -0.23)** | **-0.62 (-1.16, -0.06)** |
| Morph: Satellite | -0.12 (-0.37, 0.13) | 0.41 (-0.01, 0.83) | 0.29 (-0.07, 0.64) | **0.75 (0.16, 1.34)** |
| Morph: Faeder | 0.26 (-0.04, 0.56) | 0.47 (-0.07, 0.96) | -0.08 (-0.54, 0.38) | 0.21 (-0.53, 0.95) |
| Trial: First | **0.5 (0.28, 0.72)** | **-1.12 (-1.33, -0.9)** | **-0.55 (-0.74, -0.36)** | **-0.37 (-0.59, -0.13)** |
| Trial: Third | 0.17 (-0.06, 0.41) | **0.69 (0.48, 0.91)** | 0.01 (-0.19, 0.2) | 0 (-0.22, 0.23) |
| **Random effects** | | | | |
| Cohort (3 levels) | 0.01 (0, 0.03) | 0.2 (0.01, 0.54) | 0.02 (0, 0.06) | 0.37 (0, 1.32) |
| ID [c] | 0.08 (0.06, 0.11) | 0.74 (0.62, 0.88) | 0.57 (0.47, 0.68) | 0.56 (0.42, 0.75) |
| Residual | 0.68 (0.57, 0.8) | 0.61 (0.52, 0.71) | 0.47 (0.4, 0.54) | 0.3 (0.23, 0.38) |

[a] slope β is given for fixed effects and variance σ² for random effects. Credible intervals (CrI) not overlapping zero are highlighted in bold.

[b] Population mean for the reference of all other fixed effects, i.e., Sex: Female, Morph: Independent, Trial: Second.

[c] Early juv.-ontogeny: 103 levels, late juv.-ontogeny: 108 levels, first winter: 107 levels, second winter: 43 levels.

*Table S2: Random effect part of the output of four Linear Mixed Models (LMM) comparing the variance partitioning during the measured life stages displayed as percentage of the total variation explained by the random effects.*

| | **LMM – early juv.-ontogeny** | **LMM – late Juv.-ontogeny** | **LMM – first winter** | **LMM – second winter** |
|---|---|---|---|---|
| | Percentage of the total variance | | | |
| **Random effects** | | | | |
| Cohort (3 levels) | 1.3% | 12.9% | 1.9% | 31.1% |
| ID [a] | 10.4% | 47.7% | 53.8% | 44.3% |
| Residual [b] | 88.3% | 39.4% | 44.3% | 24.6% |

[a] interpreted as between-individual variation; early juv.-ontogeny: 103 levels, late juv.-ontogeny: 108 levels, first winter: 107 levels, second winter: 43 levels.

[b] interpreted as within-individual variation

### Fine scale repeatability development

To examine the **developmental changes in the repeatability of activity**, we calculated the adjusted individual repeatabilities (between-individual variance divided by the total phenotypic variance). Additionally, to compare the repeatability within and between the four life stages (early & late juvenile ontogeny, first & second winter), we investigated the repeatability development during the juvenile ontogeny at a finer scale by calculating the repeatability for each combination of two juvenile ontogeny trials, similar to pairwise comparisons.

The general increase of repeatability with age was also already apparent during the juvenile ontogeny trials. During the initial trials repeatability was low but then increased until reaching R = 0.6 for the final comparison between the individual trials on day 27 and 33 (Figure S4). Note that repeatability tends to be higher in this fine scale analysis, where a data subset contains only two repetitions per individual, when analysing two adjacent trials, compared to the analysis comparing the four life stages (3 trials/subset; **Fehler! Verweisquelle konnte nicht gefunden werden.** vs. Figure S4; van Berkum et al. 1989). Although it is not ideal to calculate repeatability from only two trials, as one compares the mean of two points to the mean of another two points, we see that this analysis supports the results of other models (see main manuscript, **Fehler! Verweisquelle konnte nicht gefunden werden.**).

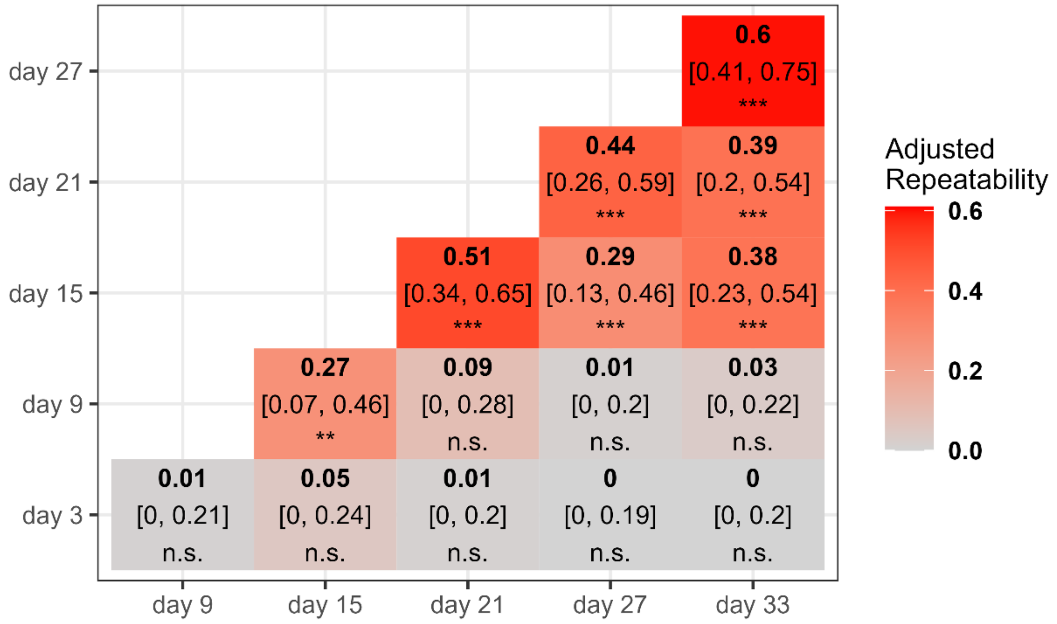


*Figure S4: Adjusted repeatabilities between juvenile ontogeny trials. The matrix contains all combinations of the single juvenile ontogeny trials. Adjusted repeatabilities are in bold, [95 % confidence interval] and significances are indicated as p-value <0.001 ***, p-value < 0.01 **, p-value > 0.05 n.s.*

### Rank stability

We investigated whether individuals establish a stable ranking within sex-morph groups based on the distance travelled throughout the juvenile ontogeny. This allowed us to assess the consistency of individual phenotypes for a trait that changes during juvenile development. High rank stability indicated that more (non)-active chicks were consistently more (non)-active relative to their peers between consecutive trials. We predicted rank stability to increase with age.

To investigate whether the chicks became more consistent in their "rank" within their sex-morph groups, we normalized the juvenile ontogeny data between 0 and 1 within each sex-morph group separately for each trial and applied a cube root transformation. Consequently, the sex, morph and age effect were eliminated and only the individual variation was left (Figure S5). Afterwards, we calculated the absolute difference between each consecutive trial for each individual (“rank”-change within sex-morph group) and applied a LMM with those absolute differences as the response variable. We included sex, morph and age as fixed effects, cohort and ID as random effects and random slopes for ID over age.

The analysis revealed that individuals became more consistent in their rank relative to peers, effectively establishing a stable ranking within sex-morph groups based on the distance travelled throughout juvenile ontogeny. Rank stability increased as the “rank”-change decreased with age. The two sexes did not differ. Satellites appeared slightly less stable than Independents and Faeders although the lower bound of the confidence interval nearly reached zero (Table S3, Figure S6).

*Table S3: Output of a Linear Mixed Models (LMM) on the individuals’ relative “ranks” within sex-morph groups during the juvenile ontogeny of activity (distance travelled in m).*

| **Response** | **Juvenile LMM - Rank**<br>“Rank”-change within sex-morph group [a] |
|---|---|
| | $\beta/\sigma^2$ (95 % CI) [b] |
| **Fixed effects** | |
| Intercept [c] | 0.20 (0.16, 0.25) |
| Sex: Male | 0.03 (-0.02, 0.07) |
| Morph: Sat | **0.06 (0.01, 0.12)** |
| Morph: Fae | 0.04 (-0.02, 0.11) |
| Age [d] | **-0.04 (-0.07, -0.01)** |
| **Random effects** | |
| Cohort (3 levels) | 0 (0, 0) |
| ID – Intercept (108 levels) | 0.01 (0.01, 0.01) |
| ID – Age [d] (108 levels) | 0 (0, 0) |
| Residual | 0.05 (0.05, 0.06) |

[a] The raw data (distance travelled in m) was first normalized between 0 and 1 within the sex-morph groups and separate for each trial, then we applied a cube root transformation and finally, we calculated the absolute differences between consecutive trials.

[b] slope β is given for fixed effects and variance σ² for random effects. Credible intervals (CrI) not overlapping zero are marked in bold.

[c] Population mean for the reference of all other fixed effects, i.e., Sex: Female, Morph: Independent, for the average age.

[d] Scaled/cantered around the mean

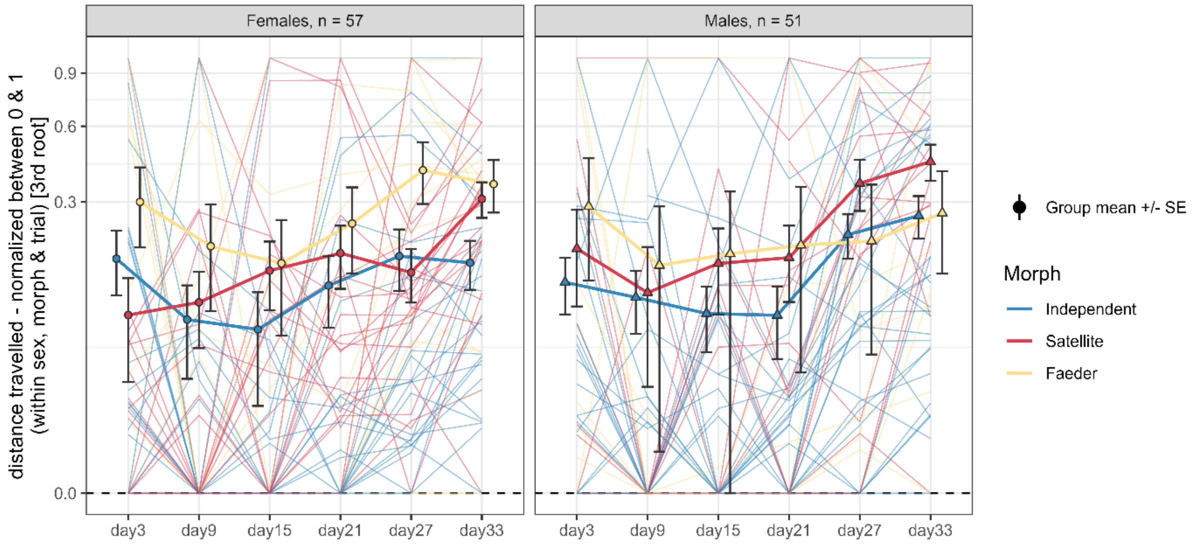


*Figure S5: Individual variation in activity (distance travelled) normalized between 0 and 1 within each sex-morph group separately for each trial. All values are displayed with cube root transformation. Consequently, sex, morph, and age effects are eliminated. The group means are plotted as symbols with standard error (error bars) connected by thick lines for females in circles (left panel) and males in triangles (right panel). Morphs can be differentiated by colour (Independents in blue, Satellites in red and Faeders in yellow). The thinner lines in the background connect all measurements of each individual.*

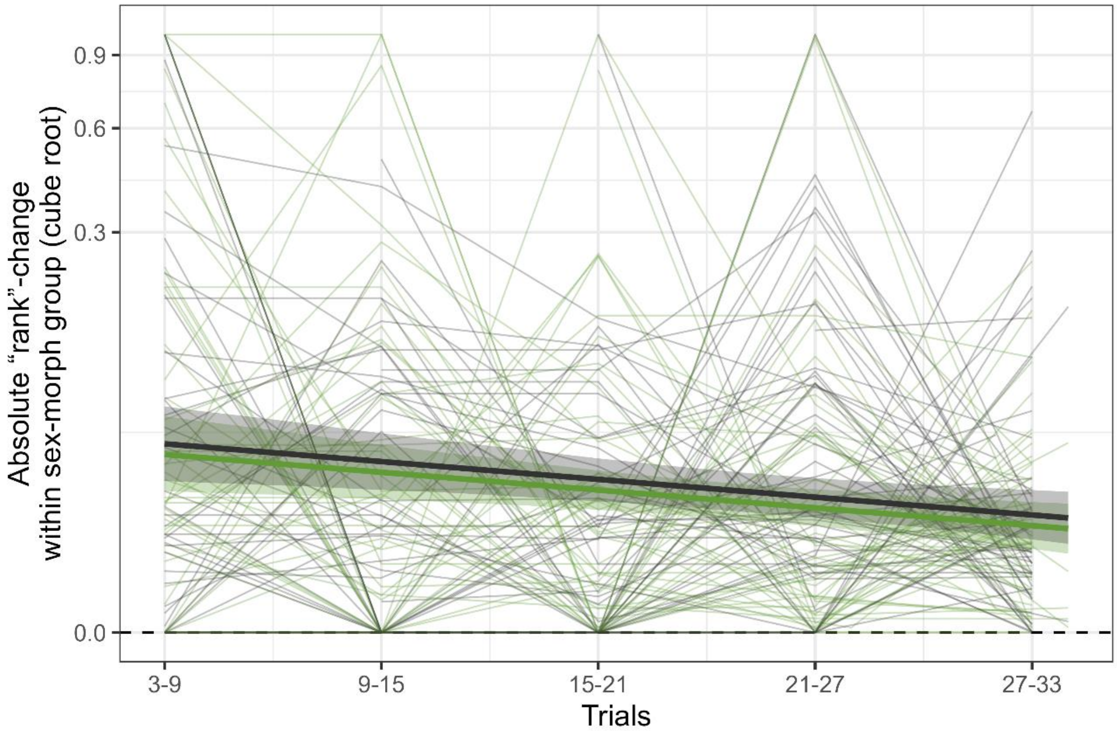


*Figure S6: Establishment of a relative "rank"-system between individuals within sex-morph groups over the juvenile ontogeny. The activity (distance travelled in m) was normalised between 0 and 1 with the sex-morph groups and separate for each trial, which removes the sex, morph and age effect and leaves only the between-individual variation. Displayed are the absolute differences between consecutive trials of these normalised values after cube root transformation. The predicted curves of the Linear Mixed Model (Juvenile LMM - Rank) are plotted as thick lines with 95 % credible intervals (shaded areas) for females in green and males in black. The thinner lines in the background connect all measurements of each individual.*

**Predictive ability**

To investigate the development of the mature phenotype of activity, we examined from which age on the juvenile behaviour can predict the mature phenotype of activity measured in the first winter. We predicted an increase of predictive ability with age, indicating more consistent behavioural responses in older juveniles.

Similar to the Winter LMM, we always used the cube-root transformed distance travelled as the response in a subset containing only the first winter data, and cohort and ID (107 levels) as random effects. As fixed effects, each of the six models included sex, morph and trial. Additionally, the distance travelled (cube root transformed and centred around the mean) during one of the six juvenile ontogeny trials (day 3, 9, 15, 21, 27 & 33) featured as a continuous predictor.

The chicks' activity during juvenile ontogeny predicted the activity observed in their first winter increasingly well. From an individual's behaviour on day 3 it is not possible to predict the activity in the first winter. However, from day 9 onwards the ability to predict an individual's later activity increases until reaching the strongest relationships on day 27 and day 33 (Table S4, Figure S7).

A similar predictive ability and consistency across life stages and different behaviours have been identified in only approximately 16 % of studies on the still understudied developmental perspective of animal personality (reviewed by Cabrera et al. 2021). Our study to our knowledge is the first to investigate the full juvenile ontogeny of activity, a personality trait, in precocial birds, alongside its long-term state. However, research covering the same period in other species also remains scarce (Cabrera et al. 2021), especially with such fine temporal resolution (but see Laskowski et al. 2022).

*Table S4: Output of four Linear Mixed Models (LMM) comparing the predictive ability of each juvenile ontogeny trials on the activity (distance travelled) measured in 450 videos in 107 individuals during the first winter.*

| | **LMM day3** | **LMM day9** | **LMM day15** | **LMM day21** | **LMM day27** | **LMM day33** |
|---|---|---|---|---|---|---|
| **Response** | Distance Travelled in the first winter (m, cube root) | | | | | |
| | $\beta/\sigma^2$ (95 % CrI) [a] | | | | | |
| **Fixed effects** | | | | | | |
| Intercept [b] | 3.42 (3.08, 3.76) | 3.45 (3.12, 3.79) | 3.44 (3.08, 3.79) | 3.43 (3.12, 3.76) | 3.45 (3.15, 3.78) | 3.4 (3.1, 3.71) |
| Sex: Male | **-0.67 (-1.03, -0.31)** | **-0.66 (-0.98, -0.32)** | **-0.59 (-0.91, -0.26)** | **-0.47 (-0.82, -0.14)** | **-0.41 (-0.72, -0.11)** | **-0.42 (-0.74, -0.09)** |
| Morph: Satellite | 0.25 (-0.15, 0.68) | 0.24 (-0.13, 0.61) | 0.21 (-0.14, 0.56) | 0.19 (-0.17, 0.56) | 0.12 (-0.23, 0.46) | 0.16 (-0.2, 0.51) |
| Morph: Faeder | -0.14 (-0.65, 0.36) | -0.19 (-0.71, 0.3) | -0.03 (-0.54, 0.45) | -0.26 (-0.72, 0.16) | -0.31 (-0.76, 0.16) | -0.18 (-0.62, 0.25) |
| Juvenile trial (m, cube root) [c] | 0.13 (-0.06, 0.31) | **0.17 (0, 0.34)** | **0.25 (0.1, 0.42)** | **0.26 (0.1, 0.43)** | **0.33 (0.17, 0.49)** | **0.3 (0.14, 0.46)** |
| Trial: First | **-0.57 (-0.77, -0.38)** | **-0.58 (-0.77, -0.39)** | **-0.57 (-0.76, -0.38)** | **-0.56 (-0.75, -0.37)** | **-0.57 (-0.75, -0.39)** | **-0.55 (-0.74, -0.36)** |
| Trial: Third | 0 (-0.19, 0.19) | 0.02 (-0.17, 0.21) | 0.02 (-0.18, 0.2) | 0 (-0.19, 0.19) | 0.01 (-0.18, 0.21) | 0 (-0.19, 0.19) |
| **Random effects** | | | | | | |
| Cohort (3 levels) | 0 (0, 0) | 0 (0, 0.02) | 0.03 (0, 0.08) | 0 (0, 0) | 0.01 (0, 0.02) | 0 (0, 0) |
| ID (107 levels) | 0.63 (0.51, 0.76) | 0.57 (0.46, 0.69) | 0.51 (0.41, 0.62) | 0.51 (0.42, 0.62) | 0.46 (0.37, 0.56) | 0.51 (0.42, 0.62) |
| Residual | 0.44 (0.37, 0.52) | 0.47 (0.4, 0.56) | 0.47 (0.4, 0.56) | 0.49 (0.41, 0.58) | 0.48 (0.41, 0.57) | 0.47 (0.4, 0.55) |

[a] slope β is given for fixed effects and variance σ² for random effects. Credible intervals (CrI) not overlapping zero with are highlighted in bold.

[b] Population mean for the reference of all other fixed effects, i.e. Sex: Female, Morph: Independent, average value measured in the ontogeny trial and Trial: Second.

[c] The juvenile ontogeny trial used as a predictor is stated in the model name (first row).

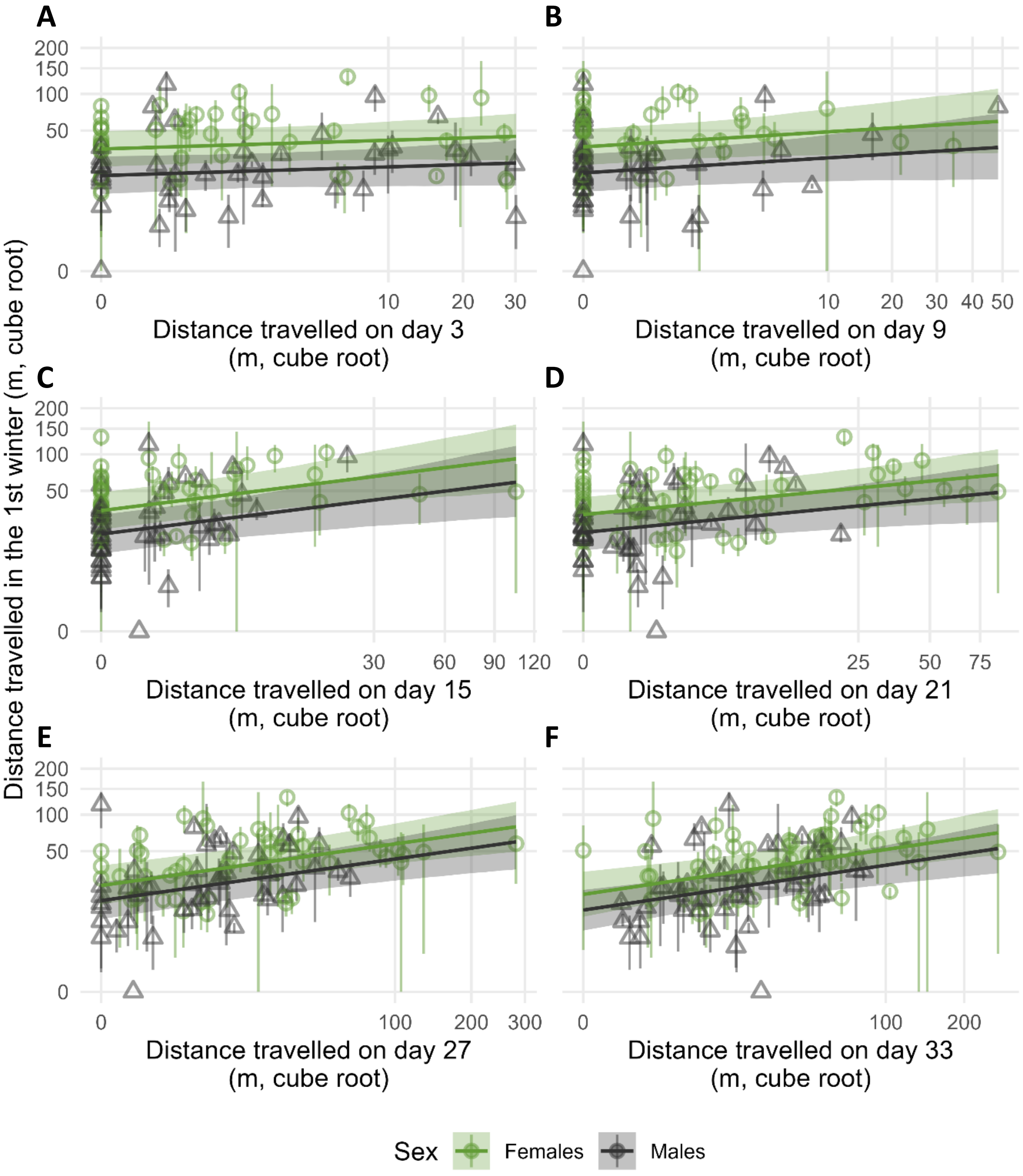


*Figure S7: Different time points during the ontogeny as predictors of the activity in the first winter. The tested time points are (A) day 3, (B) day 9, (C) day 15, (D) day21, (E) day 27 and (F) day 33. Displayed is the predicted relationship with 95 % credible intervals (shaded area). The raw data are plotted as semi-transparent point ranges indicating the minimum, mean and maximum values measured for each individual in the three first winter trials.*
